# pmparser and PMDB: resources for large-scale, open studies of the biomedical literature

**DOI:** 10.1101/2020.09.07.285924

**Authors:** Joshua L. Schoenbachler, Jacob J. Hughey

## Abstract

PubMed is an invaluable resource for the biomedical community. Although PubMed is freely available, the existing API is not designed for large-scale analyses and the XML structure of the underlying data is inconvenient for complex queries. We developed an R package called pmparser to convert the data in PubMed to a relational database. Our implementation of the database, called PMDB, currently contains data on over 31 million PubMed Identifiers (PMIDs) and is updated regularly. Together, pmparser and PMDB can enable large-scale, reproducible, and transparent analyses of the biomedical literature. pmparser is licensed under GPL-2 and available at https://pmparser.hugheylab.org. PMDB is stored in PostgreSQL and compressed dumps are available on Zenodo (https://doi.org/10.5281/zenodo.4008109).

## Introduction

As biomedical researchers continue to push into the unknown, the literature continues to grow. This growth, along with advances in technology, has created exciting opportunities at two levels. First, it enables biomedical discovery using techniques such as natural language processing (Kveler et al., 2018). Second, it enables meta-research into how research is organized, performed, disseminated, and ultimately used (Boyack et al., 2011; Piwowar et al., 2018; Wu, Wang & Evans, 2019; Abdill & Blekhman, 2019; Hutchins et al., 2019b; Fu & Hughey, 2019).

The definitive resource for the biomedical literature is PubMed/MEDLINE, maintained by the National Library of Medicine (NLM) of the U.S. National Institutes of Health. In addition to the PubMed website, all data from PubMed are freely available through the E-utilities API and to download in bulk. However, the API is designed for specific queries or small-to moderate-scale studies, and the downloadable files store data in deeply nested XML that must first be parsed. These limitations hinder large-scale analyses of PubMed data. Other large databases such as Scopus and Web of Science are not freely available, which limits access to researchers at particular institutions and discourages reproducibility and transparency.

To address these issues, we developed two companion resources: pmparser and PMDB. pmparser is an R package allowing one to easily create and update a relational database of the data in PubMed. PMDB is our implementation of the database, which is publicly available and updated regularly.

## Materials and Methods

The pmparser R package relies on the xml2, data.table, and DBI packages, which provide efficient parsing of XML documents into tables, manipulating the tables, sending them to a database, and manipulating the database from R. pmparser supports four database management systems: PostgreSQL, MariaDB, MySQL, and SQLite. The last is recommended only for small-scale testing.

NLM releases PubMed/MEDLINE data as a set of baseline XML files each December. Updates to the baseline, also XML files, are released daily (called post-baseline files below). Each file typically contains data on tens of thousands of PMIDs. pmparser parses the data into a set of tables organized by data type and linked by PMID (Table S1).

### Creating the database

To create the database, pmparser does the following:

1. Download the baseline XML files.
2. Initialize the tables in the database.
3. For each baseline file (in parallel) and for each data type:

a. Parse the XML into R data.table(s).
b. Append the data.table(s) to the corresponding table(s) in the database.
4. Add rows to the table containing the date and time at which each file was processed.
5. For each table in the database, keep only rows corresponding to the latest version of each PMID. The vast majority of PMIDs have only one version, even when their data is subsequently updated. Exceptions are articles from journals such as F1000Research.
6. Download the latest version of the NIH Open Citation Collection (Hutchins et al., 2019a) and add it as a table in the database.

### Updating the database

The post-baseline XML files contain data on new, updated, and deleted PMIDs. The procedure here is similar to above, except the data are first parsed into a set of temporary tables before being appended to the main tables.

1. Download the post-baseline XML files that have not yet been processed.
2. Initialize the temporary tables in the database.
3. For each post-baseline file (in parallel) and for each data type:

a. Parse the XML into R data.table(s).
b. Append the data.table(s) to the corresponding temporary table(s) in the database.
4. Add rows to the table containing the date and time at which each file was processed.
5. Determine the set of PMIDs whose data have been updated, along with each PMID’s most recent version and the most recent XML file in which it was updated.
6. For each main-temporary table pair:

a. Delete rows in the main table for PMIDs whose data have been updated.
b. Append rows in the temporary table that contain each PMID’s most recent data (based on version number and XML file). For instance, if a PMID is updated once in one XML file and again in a subsequent XML file, only the rows from the second update are appended. If a PMID is marked as deleted in an XML file, then that PMID’s rows are deleted from the main table, but no rows are appended.
7. Delete the temporary tables.
8. If a newer version of the NIH Open Citation Collection exists, download it and overwrite the corresponding table in the database.

## Results

Using pmparser, we created our implementation of the database, PMDB, from the baseline XML files and the latest version of the NIH Open Citation Collection. Creating the database using 24 cores on a computer with 128 GiB of memory took 4.9 hours. Updating the database from the 398 post-baseline XML files released between December 15, 2019 and October 2, 2020 took another 6.7 hours. PMDB is stored in PostgreSQL. The most recent compressed dump is 20.7 GiB and includes data for 31,601,137 PMIDs (Table S2). We plan to update PMDB monthly.

As examples of PMDB’s utility and scale, we quantified the number of authors per published article over time (Fig. 1) and the fraction of authors with an ORCID identifier over time (Fig. 2). Code to reproduce these results is available on Zenodo (https://doi.org/10.5281/zenodo.4004909). Because the queries involve multiple joins and millions of PMIDs, performing these analyses using the E-utilities API would be difficult and time-consuming. In contrast, using PMDB makes them straightforward and fast (a few minutes on our local instance).

**Figure 1.**
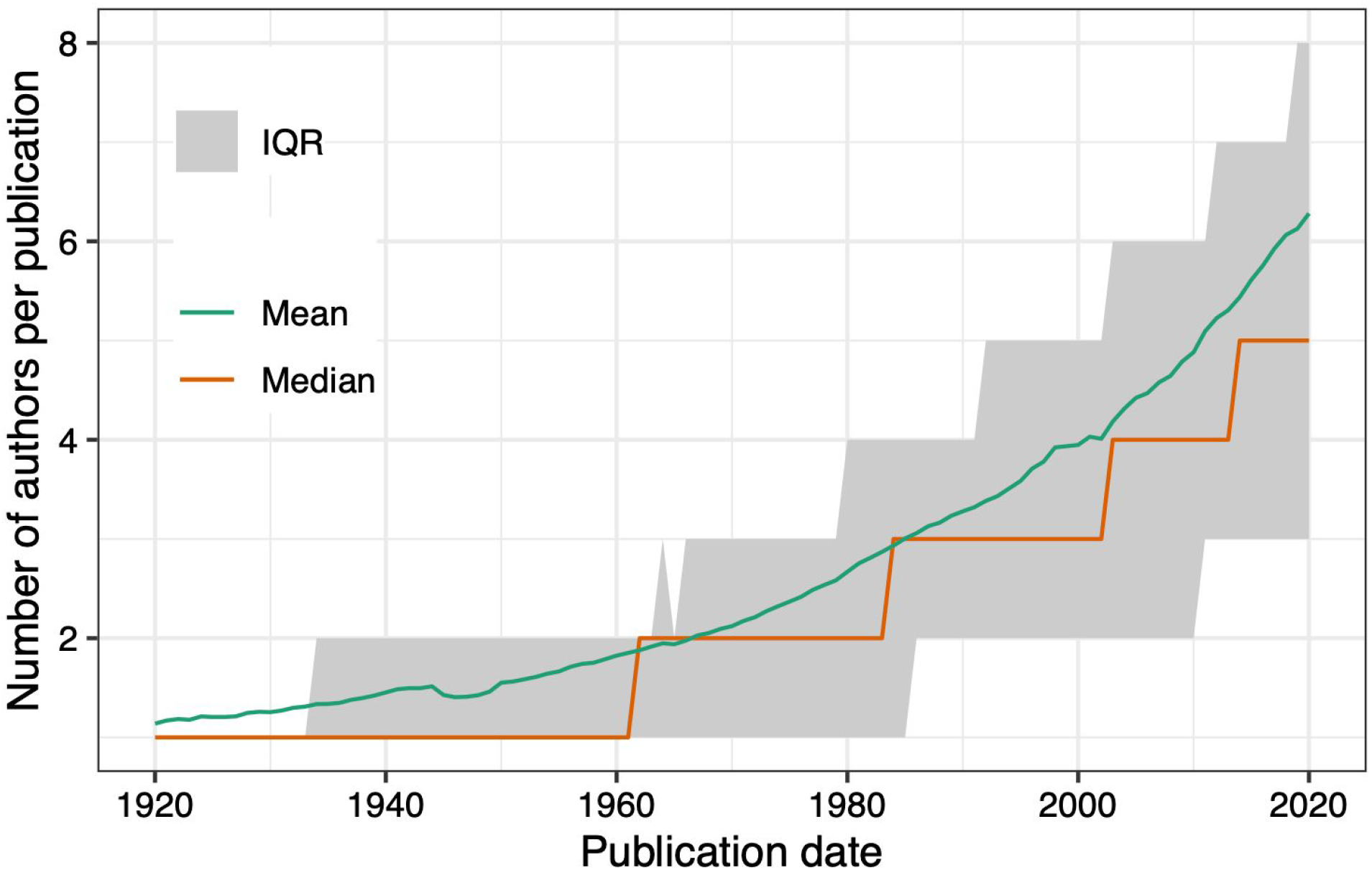
Summary statistics of the number of authors per publication between 1920 and 2020, grouped by year. The MEDLINE XML documentation states that for PMIDs created between 1984 and 1995, at most 10 authors were entered, and for PMIDs between 1996 and 1999, at most 25 authors were entered.

**Figure 2.**
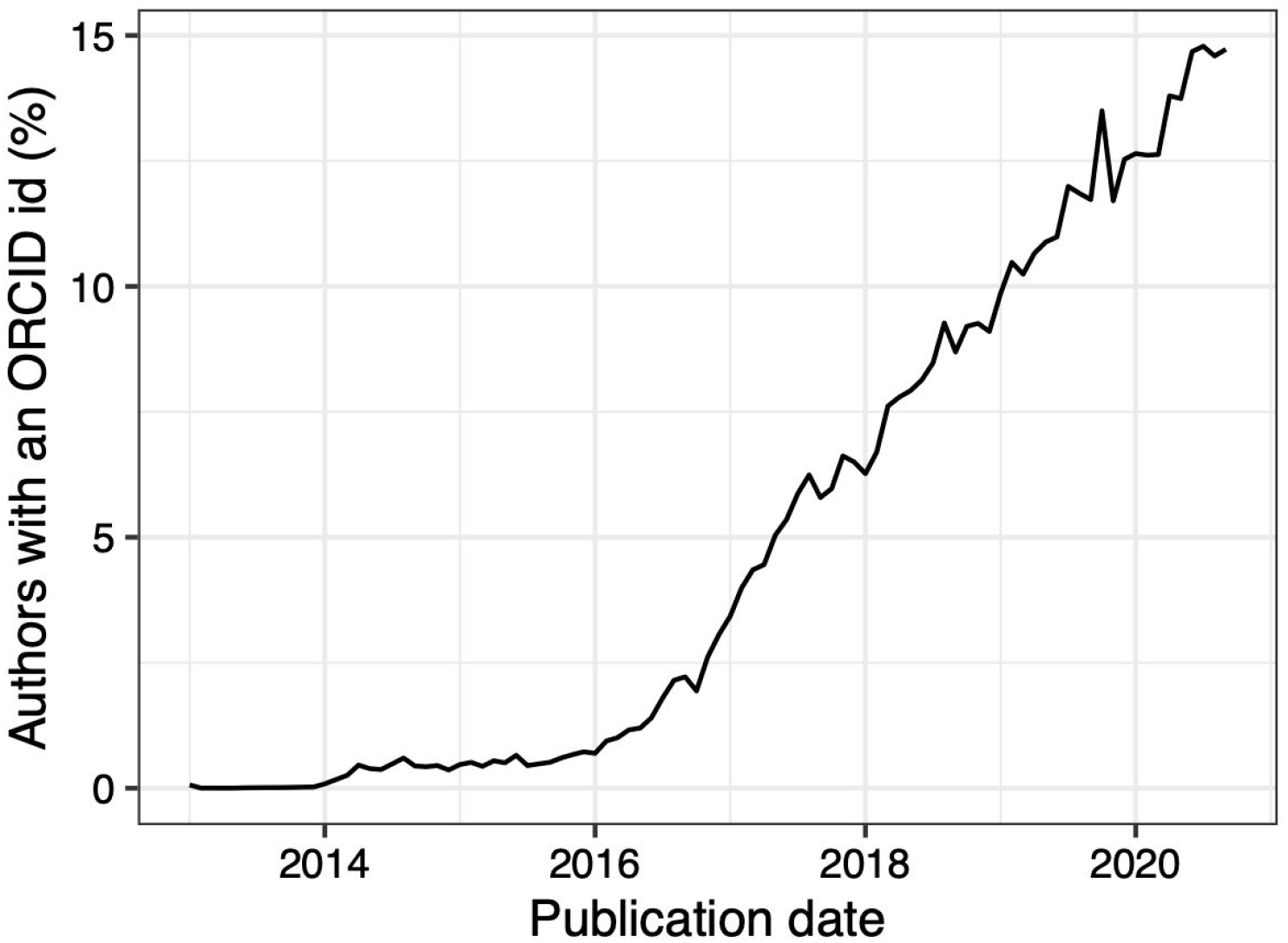
Percentage of authors with an ORCID identifier from January 2013 through August 2020, grouped by month. ORCID identifiers became available in October 2012.

## Discussion

pmparser offers similar functionality to a recently developed Python package called pubmed_parser (Achakulvisut, Acuna & Kording, 2020), with several differences. First, the packages differ somewhat in which elements of the XML they parse. For example, pmparser parses the History section, which contains data on when an article was submitted, accepted, etc., but pubmed_parser does not. Second, although both packages parse the abstracts (a common data source for biomedical NLP), pubmed_parser can also parse the full text of articles in the PubMed Central Open Access Subset. Third, pmparser adds each XML file’s data to tables in a relational database, whereas pubmed_parser outputs the data as a list of Python dictionaries. Finally, pmparser can update an existing database, whereas pubmed_parser is designed to process an entire corpus at once. Overall, which package is more suitable will depend on the use case.

With pmparser and PMDB, researchers seeking to use the wealth of data in PubMed have two new options: create their own implementation of the database or duplicate ours. Either way, once the database is set up, users can query it directly or pull the data into their tool of choice for statistical analysis, machine learning, etc. Together, pmparser and PMDB can enable large-scale, reproducible, and transparent analyses of the biomedical literature.

## Acknowledgments

We thank Elliot Outland for contributing to pmparser. We thank Darwin Fu for helpful comments on the manuscript.

